# Augmenting biologging with supervised machine learning to study *in situ* behavior of the medusa *Chrysaora fuscescens*

**DOI:** 10.1101/657684

**Authors:** Clara Fannjiang, T. Aran Mooney, Seth Cones, David Mann, K. Alex Shorter, Kakani Katija

## Abstract

Zooplankton occupy critical roles in marine ecosystems, yet their fine-scale behavior remains poorly understood due to the difficulty of studying individuals *in situ*. Here we combine biologging with supervised machine learning (ML) to demonstrate a pipeline for studying *in situ* behavior of larger zooplankton such as jellyfish. We deployed the ITAG, a biologging package with high-resolution motion sensors designed for soft-bodied invertebrates, on 8 *Chrysaora fuscescens* in Monterey Bay, using the tether method for retrieval. Using simultaneous video footage of the tagged jellyfish, we develop ML methods to 1) identify periods of tag data corrupted by the tether method, which may have compromised prior research findings, and 2) classify jellyfish behaviors. Our tools yield characterizations of fine-scale jellyfish activity and orientation over long durations, and provide evidence that developing behavioral classifiers on *in situ* rather than laboratory data is essential.

**Summary Statement:** High-resolution motion sensors paired with supervised machine learning can be used to infer fine-scale *in situ* behavior of zooplankton for long durations.

## Introduction

As anthropogenic impacts continue to alter the oceans, understanding the role movement and behavior play in how marine animals respond is required for effective stewardship and conservation. Researchers have made great strides in investigating marine megafauna behavior related to long distance migrations (Block et al., 2011; Rasmussen et al., 2007; Sequeira et al., 2018) and foraging strategies (Sims et al., 2008; Weise et al., 2010). However, the behavior of more numerous, higher total-biomass, lower trophic-level animals like zooplankton is much less understood. Early attempts to investigate *in situ* behavior of zooplankton such as jellyfish relied on scuba divers following animals with hand-held video cameras (Colin and Costello, 2002; Costello et al., 1998) and later with remotely operated vehicles (ROVs; Kaartvedt et al., 2015; Purcell, 2009; Rife and Rock, 2003). Acoustic methods have also been used to describe large-scale movement patterns of jellyfish (Båmstedt et al., 2003; Kaartvedt et al., 2007; Klevjer et al., 2009); however, these methods can be resolution-limited.

A promising alternative is biologging, where electronic transmitters or loggers with environmental and motion sensors are affixed to organisms (Kooyman, 2004; Rutz and Hays, 2009). Biologging has enabled a diverse array of marine vertebrate studies (Block et al., 2011; Goldbogen et al., 2006; Johnson and Tyack, 2003), while several technological challenges have hindered the widespread use of biologging to study gelatinous invertebrates like jellyfish. Their sensitivity to drag induces constraints on tag size, shape, and buoyancy (Fossette et al., 2016; Mills, 1984; Mooney, Katija, Shorter et al., 2015), which, coupled with bandlimited transmission capabilities, often restricts sensor payloads to low-resolution depth or location pingers (Honda et al., 2009; Moriarty et al., 2012; Seymour et al., 2004). As a result, very few studies have successfully deployed high-resolution motion sensors like accelerometers on jellyfish *in situ* (Fossette et al., 2015), and these adopted the “tether method” for retrieval (Fossette et al., 2016; Hays et al., 2008), where the tag is tethered to a surface float transmitting location. As tethering can restrict movement, it is unknown whether data collected in this manner is broadly representative of natural behavior. Furthermore, without validation from simultaneous observation of the tagged animal, interpretation of biologging data is easily biased (Brown et al., 2013; Jeantet et al., 2018).

Recently, techniques from supervised machine learning, which automatically fit or “learn” patterns that optimally distinguish categories, have been successfully used to classify behaviors in various marine vertebrates (Brewster et al., 2018; Jeantet et al., 2018; Ladds et al., 2016). However, few studies develop their methods on ground-truthed *in situ* data, due to the difficulty of recording sustained observations of wild marine animals (Carroll et al., 2014). It is unknown whether classifiers developed on data from captive, controlled, or laboratory conditions are equally effective on data from natural environments (Carroll et al., 2014), a broader problem known as domain adaptation in machine learning (Pan and Yang, 2010; Zhang et al., 2013).

In this study, we demonstrate how to investigate fine-scale zooplankton behavior *in situ* by combining biologging advancements with supervised machine learning (ML) methods. We study the movements of the scyphomedusa *Chrysaora fuscescens* in Monterey Bay, CA, USA, using the ITAG, a biologging tag equipped with high-resolution motion sensors and engineered specifically for soft-bodied invertebrates (Mooney, Katija, Shorter et al., 2015). We use the tether method for retrieval and simultaneously record video footage of the tagged animals. We develop classifiers using the resulting data to 1) detect when the tether method influences jellyfish behavior, and 2) distinguish swimming from drifting. We provide principled estimates of the classifier error characteristics, which allow us to remove behavioral data influenced by tethering, and estimate the fine-scale *in situ* orientation and swimming activity of *C. fuscescens* individuals for up to 10 h. By combining a highly specialized tag with supervised ML, our approach is the first complete pipeline for acquiring and interpreting high-resolution motion data from individual jellyfish or other zooplankton *in situ*.

## Methods & Materials

### Laboratory Deployments

Laboratory investigations of jellyfish tagging were conducted at the Monterey Bay Aquarium Research Institute (MBARI) in Moss Landing, CA in the spring of 2018. Four jellyfish (*Chrysaora fuscescens*) with bell diameters ranging from 16 to 25 cm were collected in Monterey Bay from R/V Paragon (CADFW permit SC-13337) and kept in plastic bags filled with unfiltered seawater. Within 4 hours of collection, animals were transported into large holding tanks in a 5° C cold room in MBARI’s Seawater Lab. Experiments were conducted in MBARI’s Test Tank, a 275,000 gallon tank with dimensions of 13 m (L) × 10 m (W) × 10 m (D). Animals were transported from the Seawater Lab in plastic bags and placed in the Test Tank to acclimate for at least an hour prior to tagging trials. After acclimation, a neutrally buoyant bio-logging tag (ITAG v0.4; Mooney, Katija, Shorter et al., 2015) was prepared for attachment. The ITAG (6.3 cm × 2.9 cm × 1.6 cm, air weight 30 g) is equipped with a triaxial accelerometer, gyroscope, and magnetometer synchronously sampling at a rate of 100 Hz (TDK Invensense MPU9250, San Jose, CA, USA), and pressure, temperature (TE Connectivity MS5803, Schauffhausen, Switzerland), and light sensors (Intersil ISL29125, Milpitas, CA, USA) sampling at 1 Hz. The tag was attached to the animal’s aboral surface using veterinary-grade tissue adhesive (3M Vetbond, Maplewood, Minnesota, USA), following the “glue method” (Fossette et al., 2016). Care was taken to center the attachment site on the bell apex between the four gonads, so that the tag axis conventions aligned with the jellyfish, and the animal’s radial symmetry was not disrupted. The entire attachment procedure took no longer than 2 minutes.

To replicate the *in situ* recovery strategy, the tags were attached by 6 m of monofilament line (20-lb. test) to a suspended walkway about 1 m above the tank surface (the tether length was set to prevent the animal from getting tangled with metal bars on the walls of the test tank). Simultaneous lateral-view video footage of the tagged jellyfish was collected with a HERO5 Black GoPro (GoPro, Inc., San Mateo, CA, USA) mounted onto a BlueROV2 (Blue Robotics, Torrance, CA, USA). Footage was synchronized with the tag data by sharply tapping the tag five times in front of the GoPro prior to attachment.

### Field Deployments

We deployed ITAGs on 8 *C. fuscescens* in Monterey Bay, CA in late spring of 2018, and collected *in situ* recordings with durations between 54 min and 10 h. The bell diameters of these animals were between 20 and 28 cm. Fig. 1A-G depicts the main phases of the deployment protocol. Each animal was first spotted from aboard the R/V Paragon, which then maneuvered next to the animal so it could be gently captured and brought aboard using a plastic bucket. Captured jellyfish were then transferred into individual 27-gallon plastic holding tubs filled with seawater (Fig. 1A), with care taken to not introduce air bubbles under their bells. To recover tags at the end of the deployment, we used the “tether method” (Fossette et al., 2016): tags were tethered by 30 m of monofilament line to the bottom of a drogue, which was attached by dock line to a surface drifter. A fishing swivel was placed at the midpoint of the tether, as well as immediately below its attachment to the drogue, to prevent any tether torsion from affecting the animal. The surface drifter consisted of PVC housing for a SPOT GPS tracking device (SPOT LLC, Milpitas, CA, USA), and a PVC pipe chamber containing batteries and ballasting material. The SPOT was configured to report its coordinates once every 15-20 minutes via email. The tethered tags were then affixed to jellyfish while in the holding tubs, following the aforementioned glue method also used for laboratory deployments (Fig. 1B-C). In order, the drogued drifter, ROV, and finally tagged jellyfish were then released (Fig. 1D-F, respectively). The drogue was centered at a depth down to 9 m (see Table S1), and the jellyfish could therefore swim freely down to a depth between 30 and 39 m. From pilot control on the deck, we used the ROV-mounted GoPro to track and record video footage of the tagged animal, until losing sight of it due to water turbidity and/or turbulence. Once visual contact with the tagged animal was lost, the ROV was recovered, and the tagged animals tethered to the drogued drifters were left behind.

**Figure 1.**
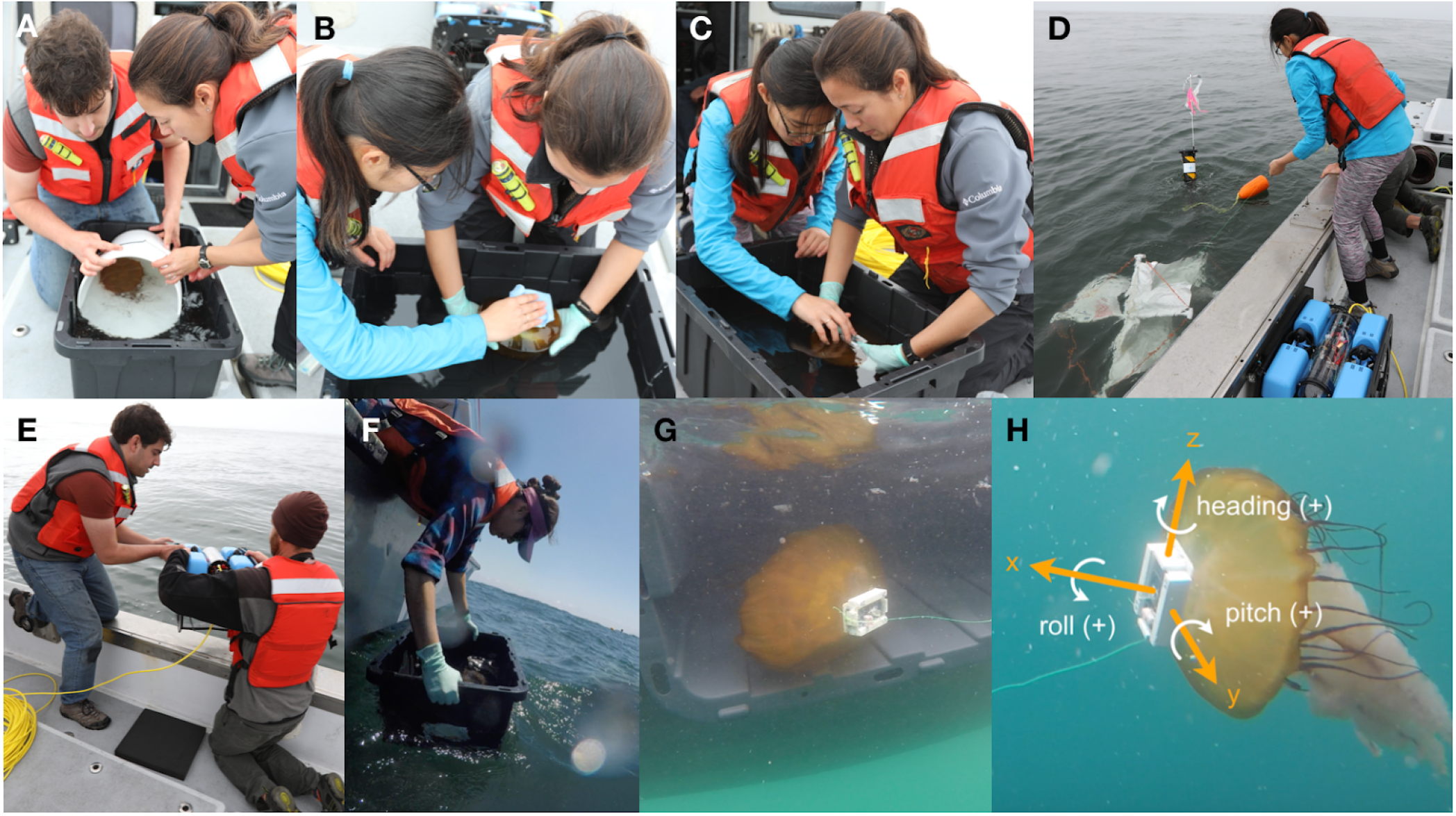
Photos of the in situ protocol and tag axis definitions. Protocol consisted of transferring collected jellyfish to staging tub (A), drying the attachment site with absorbent towels (B), gently affxing tethered ITAG with VetBond (C), deploying SPOT drifter and drogue (D), deploying BlueROV with mounted GoPro (E), and gently releasing tagged jellyfish and tracking it with the BlueROV (F-G). (H) Definitions for positive x, y, and z tag axes, and positive heading, roll, and pitch angle.

Tags were retrieved the next morning after deployment. The Paragon was navigated to the most recent coordinates reported by the SPOT, and once the drifter and drogue were located, the drifter, drogue, and tag (with or without an animal still attached) were recovered. Data from the tag was then brought back to shore for analysis.

### Orientation Estimation

We defined axes conventions appropriate for the typical jellyfish swimming position (see Fig. 1H), according to which the ITAG pressure sensor and triaxial accelerometer, gyroscope, and magnetometer were calibrated from bench tests. Since we have control over the tag attachment site, we can ensure that the tag x-axis (surge direction) is orthogonal to the jellyfish bell at the apex, so no further data processing is necessary to align the tag and jellyfish axes.

In order to compute orientation from the accelerometer and magnetometer signals, we first used a finite impulse response filter to smooth the accelerometer and magnetometer data (Sato et al., 2003). The filter cut-off frequency was set to 0.2 Hz, within the typical range of 0.25-0.5 of the pulse frequency (Martín López et al., 2016) of about 0.6 Hz estimated for a *Chrysaora* species (Matanoski et al., 2001). Filtering the accelerometer data separates the signal due to gravity (static acceleration, SA) from high-frequency animal-generated forces (dynamic acceleration or DA; Wilson et al., 2006), which we later process and featurize for behavioral classification. The resulting SA was then combined with the smoothed magnetometer data to calculate orientation (Euler angles of heading, pitch, and roll) at every point in time, according to trigonometric relationships (Johnson and Tyack, 2003). Based on our axes conventions in Fig. 1A, heading refers to compass bearing from true north, positive pitch means the jellyfish bell apex is tilted upward with respect to the horizon, and positive roll means the jellyfish bell is rotating around its apex counterclockwise, when viewed facing the bell.

### Annotation of Video Data

Throughout this paper, we refer to tag data paired with annotated simultaneous video footage as annotated data, and tag data after video footage ended as unannotated data.

Laboratory and *in situ* video footage was manually annotated for jellyfish behavior and tether influence. Each second of footage was labeled according to whether the jellyfish was swimming or drifting (not actively pulsing its bell), and whether the tether was slack (i.e. when the animal was uninfluenced by tether tension) or taut (i.e. when the animal was influenced by tether tension). For *in situ* deployments, if the tether could not be clearly seen or was out of view due to turbidity and/or viewing angle (e.g. facing the subumbrella), the state of the tether was annotated as unknown. Similarly, if the jellyfish behavior could not be distinguished due to lighting or turbidity, the behavior was annotated as unknown. Any segments of footage where either the tether state or jellyfish behavior were unknown were excluded from training data for the methods we describe below.

### Jellyfish Behavior Classification

When using the tether method as a tag retrieval strategy, prolonged deviations between the trajectories of the jellyfish and its tethered drifter can result in the tether pulling on the tag. These forces leave measurable signatures in the motion sensor data, which are distinct from the signals generated by the jellyfish’s natural behavior. Our goal was to develop supervised machine learning methods to 1) detect and remove segments of data corrupted by tether influence (tether influence classification), and 2) distinguish swimming from drifting on the remaining data (activity classification). In the following sections, we describe how these methods were developed.

### Data Preprocessing

*In situ* data was first split into two pools. Annotated data was processed and featurized as described below, then set aside for model training and evaluation. Unannotated data was similarly processed and featurized, and then set aside for classification by the trained models. Laboratory data was completely annotated, since we were able to capture video footage of the entire deployment.

We used the following procedure to assemble data samples for each of the four categories annotated as described above: tether-influenced, uninfluenced, swimming, and drifting. Upon visual inspection, the DA of every deployment displayed a nearly constant periodic nature, consistent with the nearly constant jellyfish bell pulsing observed in both the laboratory and *in situ* video footage. We therefore computed the discrete cosine transform (DCT) of the DA and took the frequency with the maximum absolute coefficient as the representative pulse frequency (RPF) for each deployment.

For each category, we extracted all segments of motion sensor data whose corresponding video footage was annotated with that category. Each segment, which consisted of 10 channels of data (pressure sensor and triaxial accelerometer, gyroscope, and magnetometer) was then split into consecutive, non-overlapping windows with a duration equal to the representative swimming cycle length (the reciprocal of the RPF). Segments shorter than this duration, and trailing windows at the ends of segments shorter than this duration, were discarded from classification and analysis. Each of these windows, which we refer to as periods, was then featurized. Note that the period duration is different for each deployment, to account for the pulse frequency of each animal.

### Featurization

For each period, we generated a total of 46 candidate features from the accelerometer and gyroscope. During training, we used a feature selection method to select a subset with the greatest predictive power, as described below. In the following, triaxial jerk was calculated as the difference between consecutive triaxial accelerometry values, scaled by the sample rate of 100 Hz. Similarly, angular acceleration was calculated as the difference between consecutive gyroscope values, scaled by the sample rate.

We computed various features of partial dynamic body acceleration, or PDBA, the sum of the absolute values of the y- and z-axis of DA. PDBA is a variant of overall dynamic body acceleration (ODBA; Wilson et al., 2006), which is used extensively as a proxy for energetic input (Halsey et al., 2009; Wilson et al., 2006). By computing both PDBA and the absolute value of the x-axis of DA (DAx), we can separate energy expenditure in the direction of jellyfish propulsion from movements in the orthogonal plane (i.e. the x-axis from the y-z plane in Fig. 1H). To account for variation in propulsion force between individual jellyfish, for each jellyfish we divided the PDBA and DAx by their respective averages over the entire deployment. Analogous to PDBA and DAx, for the gyroscope data we considered the norm of the y- and z-axis (which we call partial vectorial angular velocity, or PVAV), and the absolute value of the x-axis (AVx).

Accelerometer-based features included the maximum, mean, and standard deviations of the following quantities: DAx, PDBA, the absolute value of the x-axis of jerk, and the norm of the y- and z-axes of jerk. Spectral features were the sparsities of DAx and PDBA spectra (the absolute value of the Fourier transform), as measured by the Gini index (Hurley and Rickard, 2009; Zonoobi et al., 2011) and the spectral energies of DAx and PDBA in 0.2-1.0 Hz (roughly the typical range of pulse frequencies) and 1-8 Hz. We also included the spectral energy of DAx over 8 Hz but excluded it for PDBA, because the two were too highly correlated, leading to numerically unstable covariance matrix inversions in our model. The remaining features were the number of peaks in the DAx and PDBA, as identified by a peak-detection method (Duarte, 2013), the correlation between the y- and z-axes of DA, and the average of the correlations between the x- and y-axes and x- and z-axes of DA.

The gyroscope-based features were completely analogous to the accelerometer-based features, substituting AVx, PVAV, and angular acceleration for DAx, PDBA, and jerk above, respectively.

We also computed the maximum normalized ODBA per period for behavioral analysis, where, similarly to PDBA and DAx, we first divided the ODBA signal by the average ODBA over the deployment to accommodate differences in propulsion strength between individual jellyfish. However, it wasn’t included as a classification feature due to redundancy with PDBA and DAx.

### Training Data

The video footage showed that the nature of tether influence was fundamentally different between *in situ* and laboratory deployments. In the test tank, the jellyfish simply turned slightly whenever it reached the end of the tether, whereas tether influence *in situ* took the form of sharp yanking or prolonged dragging on the jellyfish. Since our end goal was to detect tether influence *in situ*, and the nature of *in situ* tether influence was not replicated in laboratory footage, we only trained and evaluated the tether influence classifier on *in situ* data.

For training the tether-influence classifier, the annotated *in situ* data yielded 325 s of tether-influenced behavior and 2825 s of uninfluenced behavior across all deployments. Splitting this data into periods produced 83 and 1245 periods of influenced and uninfluenced data, respectively. For training the activity classifier, the annotated laboratory data yielded 366 s (68 periods) and 9201 s (3069 periods) of uninfluenced drifting and uninfluenced swimming behavior, respectively, and the annotated *in situ* data yielded 79 s (17 periods) and 2740 s (1228 periods) of uninfluenced drifting and uninfluenced swimming behavior, respectively. Since only 17 periods of uninfluenced *in situ* drifting were observed, we trained the activity classifier on the combined *in situ* and laboratory data (85 and 4297 periods of drifting and swimming, respectively) to sufficiently capture drifting behavior. To assess the value of incorporating *in situ* data for training, we also trained the classifier solely on the laboratory data.

### Classification Methods

#### Quadratic Discriminant Analysis

For both tether influence and activity classification, we trained a supervised machine learning method known as quadratic discriminant analysis (QDA; Hastie et al., 2009), a generalization of the classical linear discriminant analysis method introduced by Fisher (Fisher, 1936; Hastie et al., 2009). QDA models each category in feature space as a multivariate normal distribution with an individual mean and individual covariance matrix. That is, let *x* ∈ *R*^*p*^ denote the feature vector, where *p* is the number of features, and let *y* ∈ {0, 1} denote the categorical label (e.g. swimming vs. drifting for the activity classifier). For convention, we let category 1 refer to the minority (less frequent) category, i.e. uninfluenced for tether-influence classification and drifting for activity classification. The data is then modeled as

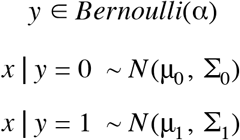

where μ_0_, μ_1_ ∈ *R*^*p*^ and Σ_0_, Σ_1_ ∈ *R*^*p*×*p*^ are the mean and covariance matrix parameters, respectively, and α ∈ [0, 1] is the probability of category 1 occurring, known as the class prior. We fit the model by computing the maximum likelihood estimates (MLE) for μ_0_, μ_1_, Σ_0_, Σ_1_, and α, which are simply the sample means and sample covariances of the categories, and the proportion of category 1 in the training set. Under this model, QDA then classifies a new instance to the category 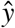 that maximizes the conditional probability 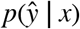 (the category that is most likely given the features), which can be accessed via Bayes’ rule. As the name implies, the resulting decision boundaries in feature space are quadratic curves. Due to the simplicity of the model and closed-form nature of the MLE, QDA is both easy to interpret and fast to train.

#### Feature Selection

There is often a large number of candidate features one can consider for a classifier. Principled methods for choosing an optimal subset of these features can help produce classifiers that perform better (due to the removal of noisy, irrelevant, or redundant features), are faster and cheaper to use (since fewer feature need to be measured and processed), and are more interpretable (Dash and Liu, 1997; Guyon and Elisseeff, 2003; Liu and Motoda, 1998). Under the broader umbrella of model selection, feature selection encourages finding the simplest model that explains the data, a principle that is critical for performance generalization (Hastie et al., 2009; MacKay, 2003). We first manually generated the list of 46 candidate features described above from accelerometer and gyroscope data. As part of training, we use a popular greedy heuristic known as sequential forward selection (SFS; Whitney, 1971), which starts with an empty subset of features and iteratively adds the next feature whose inclusion to the existing subset improves some evaluation metric the most. Despite its simplicity, SFS has been shown to match or outperform more complex search methods by being less prone to overfitting (Reunanen, 2003).

#### Metric for Feature Selection

In choosing an evaluation metric for SFS, we observed that our video annotations showed highly skewed category distributions for both classification tasks: tether-influenced periods and drifting were observed far less often than uninfluenced periods and swimming, respectively. In this case, the common metric of accuracy loses meaning, since the accuracy of a simple majority decision rule (i.e. always predict the majority categories, influenced and swimming) is high even though 1) the features are not considered and 2) all instances of the minority category are misclassified. Regardless of category imbalance, the evaluation metric should reflect how well a classifier extracts discriminating information from the features, and should account for the balance of false positives and false negatives on the minority category.

In particular, consider precision and recall on the minority category, defined as

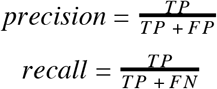

where *TP* denotes the number of true positives, or minority category periods correctly classified as the minority category; *FP* denotes the number of false positives, or majority category periods incorrectly classified as the minority category; and *FN* denotes the number of false negatives, or minority category periods incorrectly classified as the majority category. Given a trained probabilistic model of the data, such as the one posed by QDA, the decision rule to classify an instance as category 1 can be formulated in terms of a threshold on the probability *p*(*y* = 1 | *x*). Varying this threshold exposes an inherent trade-off between precision and recall: a decision rule with a high threshold, which only selects the category given overwhelmingly high evidence, tends to achieve higher precision at the cost of lower recall. A decision rule with a lenient threshold, which liberally selects the category given only mild evidence, tends to achieve higher recall at the cost of lower precision. This trade-off is captured by the curve in precision-recall space (PR curve) generated by decreasing the decision threshold from 1 to 0, which is often used to characterize classifier performance on tasks with skewed category distributions (Bunescu et al., 2005; Davis and Goadrich, 2006; Fawcett, 2006; Manning and Schütze, 1999). The PR curve allows the analyst to choose an appropriate decision threshold, depending on the relative importance of precision and recall for the task at hand (Manning and Schütze, 1999).

We use the area under this curve (AUPRC) as the metric for feature selection, which provides a summary of performance across all possible thresholds (Boyd et al., 2013; Richardson and Domingos, 2006). The AUPRC ranges from 0 to 1, where an ideal classifier that suffers no trade-off has an AUPRC of 1, and a classifier no better than random guessing has an expected AUPRC of the proportion of the category in the dataset. During feature selection, we terminate SFS when the inclusion of the next feature fails to improve the AUPRC by at least 0.02.

#### Classifier Evaluation

Unbiased evaluation of a classifier’s performance on unseen data requires complete separation of the data used in the training and evaluation phases. The standard way to evaluate classifier performance is with *k*-fold cross-validation (CV; Hastie et al., 2009; Kohavi, 1995), in which the annotated dataset is split equally into *k* parts. For each part, a classifier is trained on the remaining *k* − 1 parts (the training set) and evaluated on the excluded part (the validation set) using some evaluation metric, and the average of the resulting *k* evaluation scores (the CV score) is used as an estimate of the method’s evaluation score on unseen data. Since we want to take full advantage of our annotated dataset for training, a final classifier can then be trained on the complete dataset and deployed for future predictions (Cawley and Talbot, 2010; Varma and Simon, 2006).

Note that during evaluation, the training phase must include all aspects of model selection, including feature selection and choosing hyperparameter values. However, these aspects are sometimes incorrectly treated as external to the training process: performing hyperparameter and/or feature selection on the complete dataset, prior to CV, can result in dramatic inflations of the CV score (Ambroise and McLachlan, 2002; Cawley and Talbot, 2010; Smialowski et al., 2010; Varma and Simon, 2006). To remove this selection bias, for each of the *k* iterations of CV, we perform feature selection solely on the training set using an “inner” CV (Ambroise and McLachlan, 2002; Varma and Simon, 2006). That is, the training set is itself split evenly into *k* parts, and for each iteration of SFS, each candidate feature is evaluated by 1) adding it to the current feature subset, 2) training QDA with those features and evaluating it on the *k* pairs of training and validation sets, and 3) averaging those *k* evaluation scores. The feature with the best inner CV score is then selected. After SFS has terminated, we use the finalized feature subset to train and evaluate QDA on the outer CV training and validation set. For both the inner and outer CV, we take *k* = 5.

After the outer CV is complete, we report the average and standard error (SE) of the *k* AUPRC values. We then use the average of the *k* PR curves to choose a decision threshold (Fawcett, 2006). For the purposes of demonstrating our methods, we prioritize precision and recall equally, and simply choose the threshold value out of {0.1, 0.2, …, 0.9} that yields the closest precision and recall. We call this the equal error rate threshold (Duda et al., 2000) and report the CV precision, recall, and accuracy for this classifier. For future studies that prioritize either precision or recall over the other, the researcher can use the average PR curve to pick a threshold that achieves the desired trade-off (Manning and Schütze, 1999; Duda et al., 2000; Fawcett, 2006).

#### Activity Classifier Baselines

To see if our featurization and feature selection approach improved activity classification beyond simpler alternatives, we trained and evaluated two baseline classifiers. The first baseline, which we refer to as ODBA thresholding, simply classifies a period as swimming if the mean normalized OBDA is above some decision threshold. Since ODBA is often used as a proxy for energetic expenditure, intuition would suggest it should be sufficient for discriminating swimming from drifting in a noiseless scenario. The second baseline follows our method but only uses accelerometry features, excluding features from the gyroscope data.

#### In Situ *Behavior Prediction*

After training and evaluating the classifiers, we used them to predict tether influence and activity on the unannotated *in situ* data. After removing any periods classified as tether-influenced, we then classified each remaining uninfluenced period as swimming or drifting. These classifications provide estimates of 1) how often the tether method interferes with the natural movements of jellyfish, and 2) how much time jellyfish spend swimming versus drifting *in situ* over long durations.

### Orientation Change

To assess change in orientation during swimming, we computed the difference in heading, pitch, and roll angles between and start and end of each non-excluded period. We converted these differences into a non-negative total angle of rotation (Diebel 2006), which we refer to as orientation change over a period. We also used circular mean and circular standard deviation to compute the average and standard deviation of heading, pitch, and roll angles over periods.

To avoid ill-defined heading and roll values due to gimbal lock, here we excluded periods where the absolute pitch angle exceeded 70° from the following analysis (this removed 0.7% of total laboratory and *in situ* periods).

### Statistical Tests

We ran several statistical tests on the annotated data to investigate potential distinctions between laboratory and *in situ* behavior, and between tether-influenced and uninfluenced behavior. For the following four tests, we used the nonparametric Mann-Whitney U test to avoid any distributional assumptions on the quantities of interest, and because we expected to have a considerably large sample size (each period constitutes only a few seconds of data). Specifically, we pooled tether-influenced periods and uninfluenced periods across the *in situ* deployments, and tested whether either group tends to exhibit 1) greater normalized ODBA and 2) greater orientation change than the other. We also tested these two hypotheses between laboratory and *in situ* data, by pooling together uninfluenced periods across the laboratory deployments and across the *in situ* deployments.

## Results

### Laboratory and *In Situ* Deployments

Fig. 2A shows drifter trajectories and timestamps for the 8 *in situ* deployments in the Monterey Bay, over three separate days (see Table S1 for laboratory and *in situ* deployment details). Video footage was successfully captured for 7 of these deployments, and annotated for activity and tether influence as summarized in Table S2 (see Movie S1 for examples of annotated footage). Drifting behavior was observed in 5 deployments, and ranged from 0.5% to 4.3% of the time. Tether influence was also observed in 5 deployments (0.35% to 28.6%).

**Figure 2.**
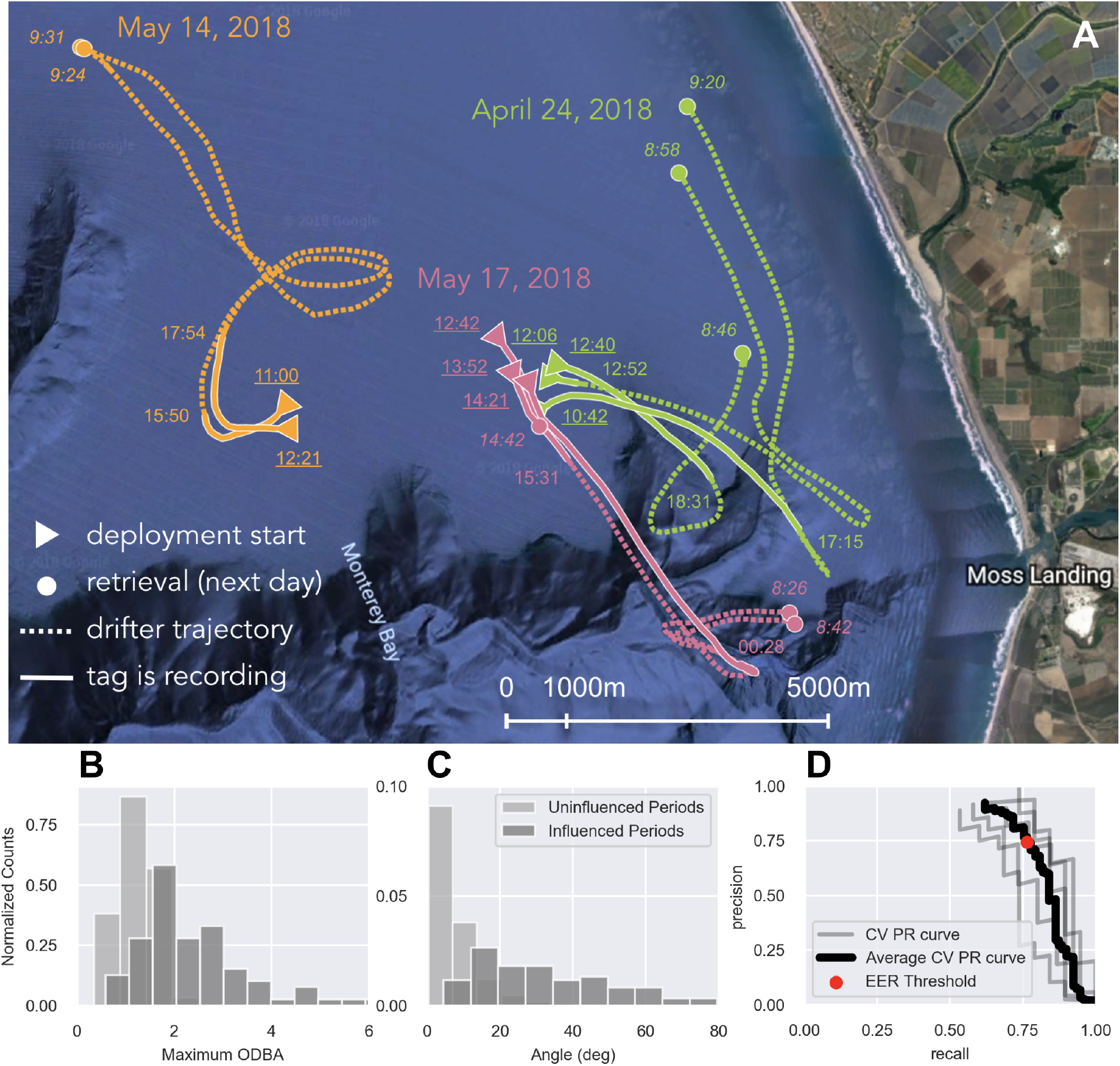
*In situ* deployment trajectories, effects of tether influence, and precision-recall curve of activity classifier. (A) Trajectories for the three deployment dates. Underlined times (PDT) denote deployment start; italicized times denote when tag was recovered; remaining times denote when tag stopped recording. (B) Maximum ODBA and (C) total orientation change over annotated tether-influenced and uninfluenced periods. (D) Cross-validation precision-recall curves of the activity classifier, and precision and recall using the equal error rate threshold.

### Jellyfish Behavior Classification

#### Jellyfish Behaviors Influenced by Tether

The tether-influence classifier had a cross-validation (CV) AUPRC of 0.860 (SE = 0.032), and using the equal error rate (EER) threshold had a CV precision of 86.1% (SE = 5.3%), recall of 73.5% (SE = 3.8%), and accuracy of 97.6% (SE = 0.4%). In order of selection by SFS, the features were 1) spectral energy of DAx over 8 Hz, 2) mean *y*-*z* PDBA, 3) number of peaks in the *y*-*z* PDBA, and 4) max DAx. After these four, SFS found no additional features that appreciably improved performance.

We used the tether-influence classifier to classify each unannotated *in situ* period as influenced or uninfluenced. Table S3 shows the proportion of each deployment classified as influenced, which ranged from 3.3% to 35.1%.

To understand how the tether influenced *in situ* behavior, we evaluated how normalized ODBA and orientation change differed between annotated uninfluenced and influenced periods (Fig. 2B, C). The maximum normalized ODBA over influenced periods (median 2.05) tended to be larger than that of uninfluenced periods (median 1.27; Mann-Whitney U test, two-sided *p* < 1e-4). Similarly, orientation change tended to be greater over influenced periods (median 30.8 degrees) than over uninfluenced periods (median 5.6 degrees; Mann-Whitney U test, two-sided *p* < 1e-4). That is, jellyfish exhibited greater ODBA and more severe orientation changes when influenced by the tether.

### Jellyfish Swimming Activity

The activity classifier had a CV AUPRC of 0.746 (SE = 0.047), and using the EER threshold had a CV precision of 74.3% (SE = 4.9%), recall of 76.4% (SE = 2.7%), and accuracy of 99.0% (SE = 0.1%). Fig. 2D demonstrates the average CV PR curve used to identify the EER threshold. The features, in order of selection by SFS, were 1) number of peaks in the PVAV, 2) sparsity of the PVAV spectrum, and 3) sparsity of the PDBA spectrum.

Note that since drifting occupied only 1.9% of the annotated periods, a simple majority prediction rule has an accuracy of 98.1%. The other metrics therefore give more insight into whether the classifier actually learns discriminative information about the categories, rather than simply which category is more common. In comparison to our method, the baseline of ODBA thresholding had an AUPRC of 0.585 (SE = 0.056) and, with the EER threshold, a precision of 68.9% (SE = 9.3%), recall of 49.9% (SE = 3.5%), and accuracy of 98.6% (SE = 0.1%). Training our classifier without gyroscope features, and only with accelerometry features, gave an AUPRC of 0.679 (SE = 0.027), and precision of 73.9% (SE = 4.6%), recall of 58.0% (SE = 7.8%), and accuracy of 98.7% (SE = 0.1%) with the EER threshold.

We used our method to classify each unannotated *in situ* period as swimming or drifting, which provided estimates of how much time each jellyfish spent for each activity. We first removed periods predicted to be tether-influenced, so that our estimates are restricted to data representative of natural behavior. The proportion of uninfluenced time each jellyfish was classified as drifting ranged between 0% and 5.6% (Table S3), with the exception of deployment S1-1 (19.1%) which also experienced frequent tether influence (both annotated and predicted). We can then combine the outputs of the influence classifier, activity classifier, and orientation estimation (again, restricted to periods predicted as uninfluenced) to visualize fine-scale information about *in situ* behavior over several hours (Fig. 3).

**Figure 3.**
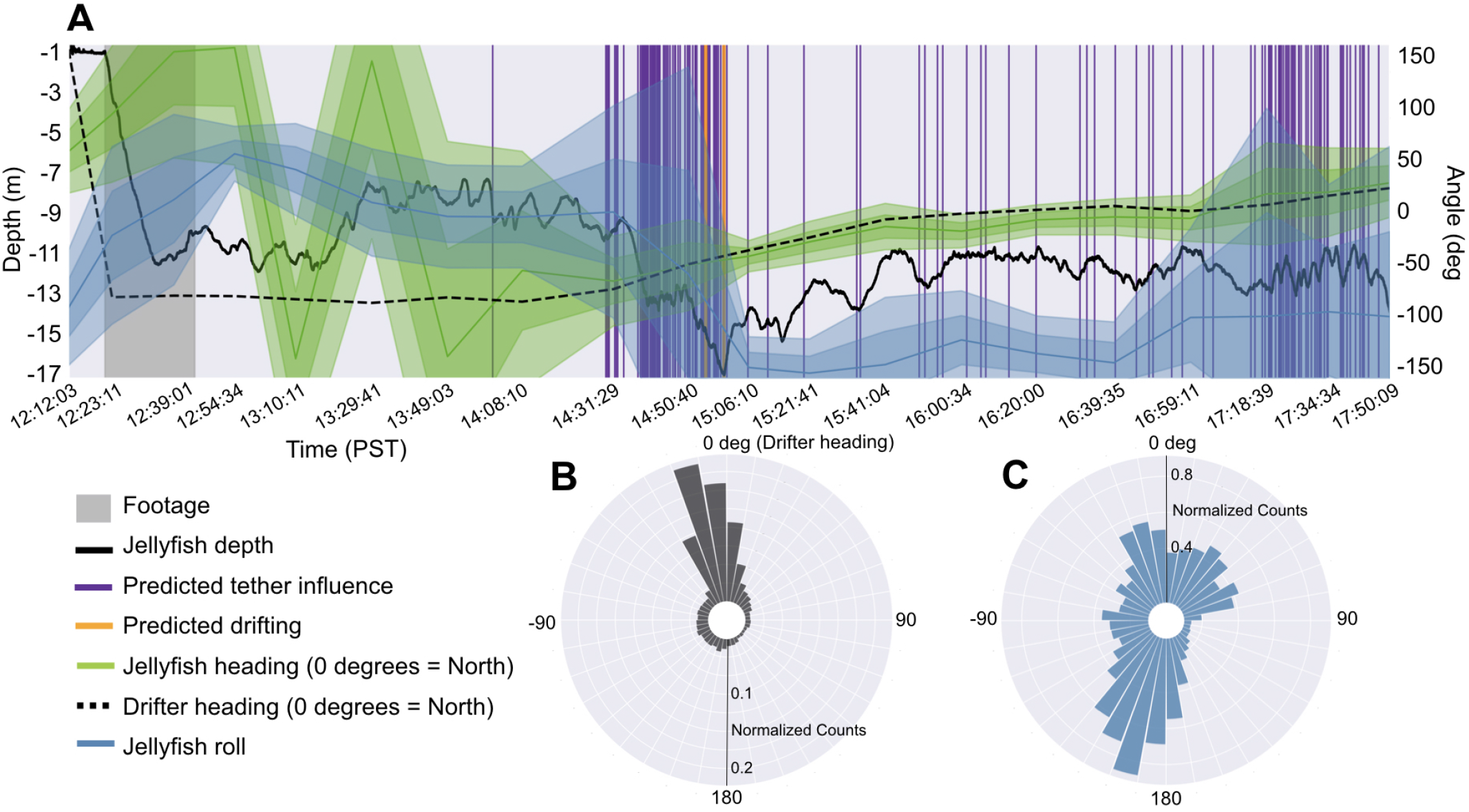
Fine-scale orientation and predicted activity of deployment S2-2. (A) Estimated heading and roll, and predicted tether influence and drifting, over entire deployment. Shaded regions denote one and two standard deviations around mean. Note that the 1-pixel-width vertical lines are disproportionately wide, as each predicted event only lasts a few seconds. (B) Radial histogram of jellyfish heading relative to the drifter heading at zero, and (C) jellyfish roll angle throughout deployment.

### Classifier Trained on *In Situ* vs. Laboratory Data

When trained and evaluated only on laboratory data, the activity classifier had a CV AUPRC of 0.894 (SE = 0.067) and, using the EER threshold, precision of 87.4% (SE = 6.1%), recall of 90.3% (SE = 4.4%), and accuracy of 99.4% (SE = 0.2%). However, predictions made by this classifier on the annotated *in situ* data had an accuracy of 96.3%, precision of 0%, and recall of 0%. We emphasize that this means none of the periods classified as drifting were truly drifting, and none of the drifting periods were correctly classified. Similarly, ODBA thresholding had an optimistic AUPRC of 0.864 (SE = 0.047), precision of 78.2% (SE = 6.3%), recall of 81.1% (SE 4.0%), and accuracy of 99.1% (SE = 0.1%) when CV was performed only on laboratory data. However, predictions on annotated *in situ* data had an accuracy of 90.8%, precision of 0%, and recall of 0%.

### *In Situ* vs. Laboratory Behavior

The maximum normalized ODBA of uninfluenced laboratory periods (median 1.98) tended to be greater than that of *in situ* uninfluenced periods (median 1.27; Mann-Whitney U test, two-sided p < 1e-4). Orientation change also tended to be greater over uninfluenced laboratory periods (median 11.6 degrees) than over uninfluenced *in situ* periods (median 5.6 degrees; Mann Whitney U test, two-sided p < 1e-4). Note that test tank walls were not responsible for turning behavior, since the tether length prevented jellyfish from reaching the walls.

## Discussion

Our work provides a pipeline for interpreting fine-scale *in situ* behavior of a zooplankton species (*Chrysaora fuscescens*) over long durations. Our approach of combining biologging with supervised ML methods yields records of *in situ* activity and orientation of individual jellyfish for several hours (up to 10 h so far), and may include the first successful *in situ* deployments of magnetometers and gyroscopes on jellyfish. Using our activity classifier, our estimates of animals’ *in situ* swimming activity on unannotated durations (on average 96.4% of the time; Table S3) is compatible with swimming in our annotated footage (on average 98.7% of behavior not annotated as unknown; Table S2). These long periods of sustained swimming with limited bouts of drifting are consistent with activity budget estimates of other oblate jellyfish (Colin et al., 2003; Costello et al., 1998), whose rowing mode of propulsion has been shown to be energy-efficient (Dabiri et al., 2010; Gemmell et al., 2018). In spite of tether influence, uninfluenced periods of data also revealed that tagged animals underwent stereotypical vertical excursions (Fig. 3A; Hays et al., 2012). Though future studies of fine-scale zooplankton behavior would be best conducted with tetherless tag retrieval methods, our approach provides a reasonably precise solution for detecting this influence and removing it, since it may compromise findings on *in situ* energetics and orientation (Fig. 2B, C; Fossette et al., 2015; Hays et al., 2008).

Our findings also highlight the importance of collecting *in situ* biologging data, rather than captive laboratory data, for developing behavioral classification methods. An assumption fundamental to justifying the deployment of machine learning (ML) methods, is that the data seen during training and inference are drawn from the same underlying distribution (Pan and Yang, 2010; Sugiyama et al., 2007; Zhang et al., 2013). Classifiers for interpreting accelerometry data, however, have been overwhelmingly trained and validated on laboratory data (Carroll et al., 2014). In doing this, these studies implicitly assume that behavioral data generated in the laboratory is distributionally similar to *in situ* behavioral data. Our findings suggest that this assumption has limited applicability, even for organisms displaying simple behaviors like swimming or drifting. First, basic descriptive statistics differed significantly between laboratory and *in situ* data: jellyfish pulses induced greater orientation changes and greater ODBA in the laboratory than *in situ*. Second, the activity classifier trained and validated solely on laboratory data had optimistic estimates of precision and recall, but performed poorly with zero precision and recall when evaluated on *in situ* data. We highlight this as a cautionary tale against naively deploying ML classifiers developed on laboratory data in the field. As biologging moves forward, methods involving technologies that capture the behavioral ground truth of *in situ* data, such as camera tags, are strongly encouraged.

Our work also underscores the limitations of ODBA in characterizing even simple *in situ* behaviors. ODBA thresholding yielded zero precision and recall in classifying *in situ* swimming and drifting, but performed reasonably well when trained and evaluated on laboratory activity. This suggests that the standard way of computing ODBA may not be robust to dynamic and unpredictable sources of noise in *in situ* data (Shepard et al., 2008). Beyond accelerometry, our results also show that leveraging information from other sensors (e.g. gyroscope) can improve *in situ* behavioral classification considerably. Looking forward, our methods open the door to investigating more complex questions about fine-scale zooplankton behavior, such as how these species orient themselves in a current, whether they exhibit rolling behavior or lateral preferences (Fig. 3), and whether their behavioral patterns distinguish them from passive drifters.

## Acknowledgments

We are grateful to Jared Figurski, Joost Daniels, Chris Wahl, William Gough, James Fahlbusch, Jeremy Goldbogen, Rob Sherlock, Steve Haddock, Chad Kecy, Tom O’Reilly, and Ivan Masmitja for guidance and assistance in the field, and to David Cade for calibration and orientation estimation tools. This work was supported by the David and Lucile Packard Foundation (to Katija), the WHOI Green Innovation Award (to Mooney, Katija, and Shorter), and NSF-DBI Collaborative awards 1455593 (to Mooney and Shorter) and 1455501 (to Katija).

## Supplementary Information

**Table S1.**
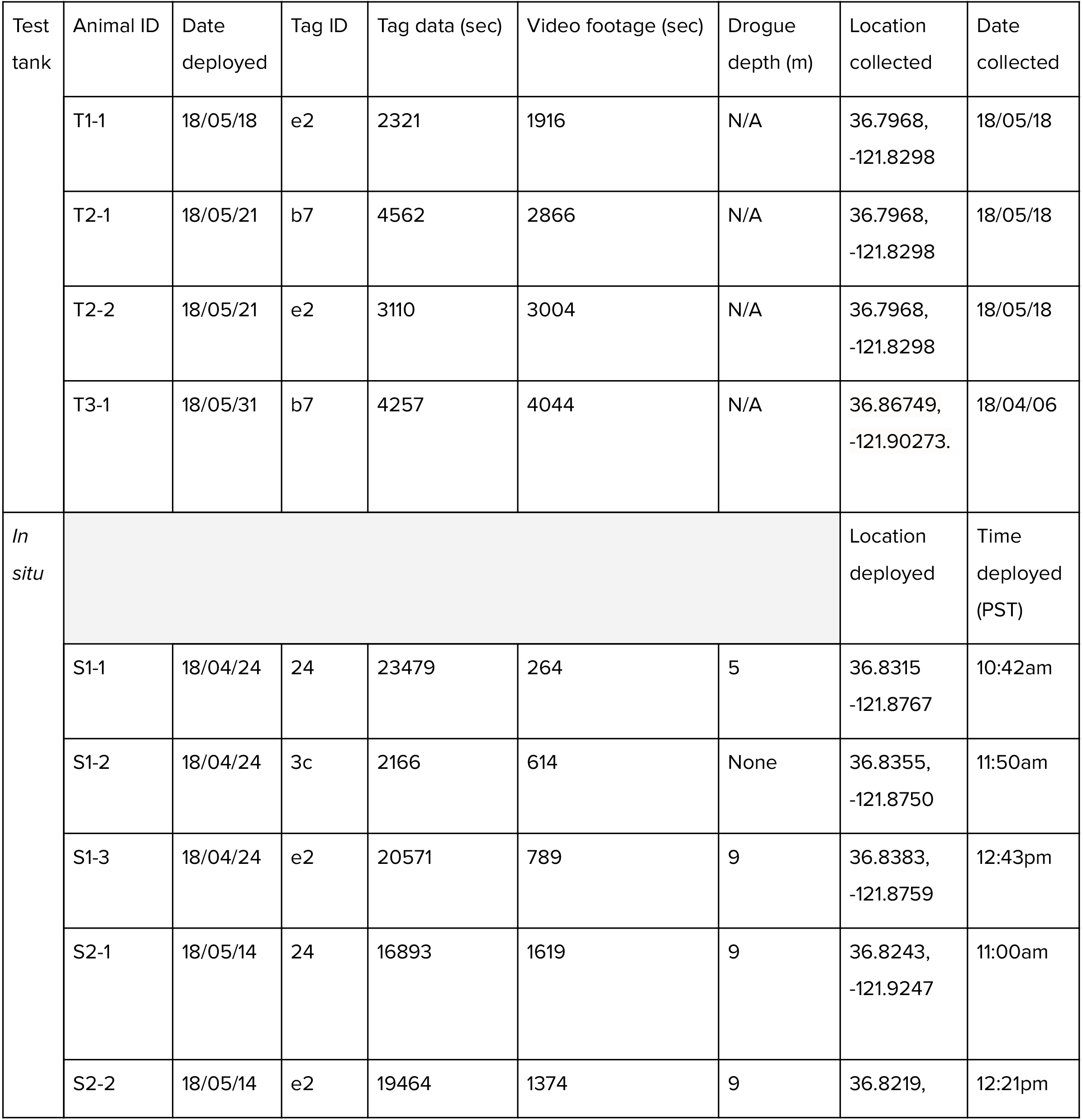
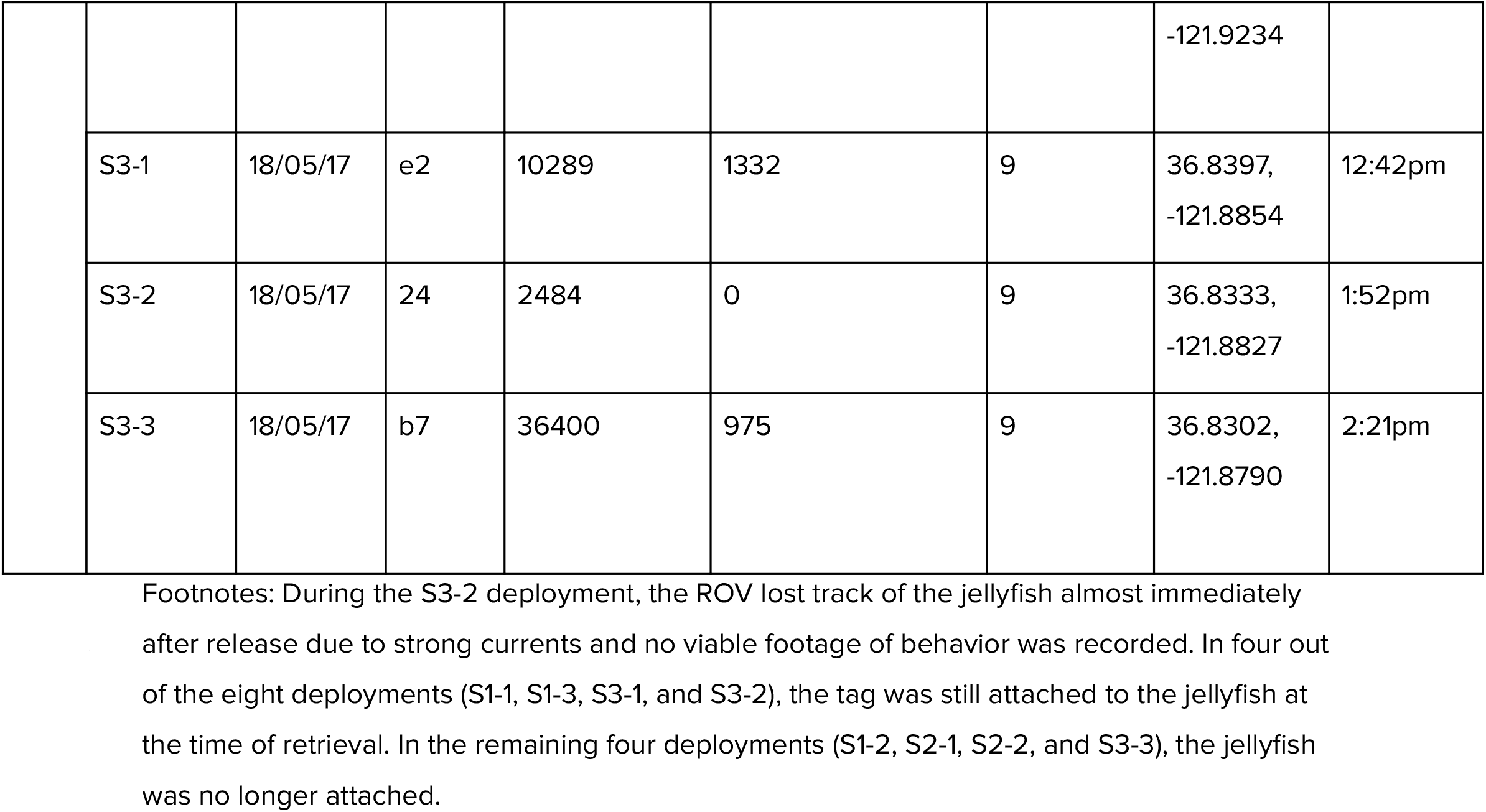
Summary of laboratory and *in situ* deployments of ITAG on *Chrysaora fuscescens*.

**Table S2.** Summary of test tank and *in situ* video footage annotations.

| Test tank | Animal ID | Total annotated footage (sec) | Behavior |  |  | Tether Influence |  |  | Unannotated tag data after footage (min) |
| --- | --- | --- | --- | --- | --- | --- | --- | --- | --- |
|  |  |  | Drift (sec) | Swim (sec) | Unknown (sec) | Taut tether (sec) | Slack tether (sec) | Unknown (sec) |  |
|  | T1-1 | 1916 | 0 | 1916 (100%) | 0 | 1093 (57.0%) | 816 (42.6%) | 7 (0.4%) | N/A |
|  | T2-1 | 2866 | 55 (1.9%) | 2811 (98.1%) | 0 | 575 (20.1%) | 2253 (78.6%) | 38 (1.3%) | N/A |
|  | T2-2 | 3004 | 4 (0.1%) | 3000 (99.9%) | 0 | 550 (18.3%) | 2454 (81.7%) | 0 | N/A |
|  | T3-1 | 4044 | 311<br>(7.7%) | 3733<br>(92.3%) | 0 | 0 | 4044 (100%) | 0 | N/A |
| Total test tank |  | 11830 | 370<br>(3.1%) | 11460<br>(96.9%) | 0 | 2218<br>(18.7%) | 9567<br>(80.9%) | 45 (0.4%) | N/A |
| *In situ* | S1-1 | 154 | 0 | 154<br>(100%) | 0 | 24 (15.6%) | 102 (66.2%) | 28 (18.2%) | 388 |
|  | S1-2 | 590 | 3<br>(0.5%) | 350<br>(59.3%) | 237<br>(40.2%) | 25 (4.2%) | 136 (23.1%) | 429 (72.7%) | 37 |
|  | S1-3 | 653 | 5<br>(0.8%) | 631<br>(96.6%) | 17 (2.6%) | 127 (19.4%) | 207 (31.7%) | 319 (48.9%) | 339 |
|  | S2-1 | 1431 | 7<br>(0.5%) | 1415<br>(98.9%) | 9 (0.6%) | 0 | 727 (50.8%) | 704 (49.2%) | 262 |
|  | S2-2 | 1347 | 0 | 1347<br>(100%) | 0 | 0 | 1285 (95.4%) | 62 (4.6%) | 311 |
|  | S3-1 | 1158 | 31<br>(2.7%) | 1116<br>(96.4%) | 11 (0.9%) | 69 (6.0%) | 285 (24.6%) | 804 (69.4%) | 146 |
|  | S3-2 | 0 | N/A | N/A | N/A | N/A | N/A | N/A | 54 |
|  | S3-3 | 762 | 33<br>(4.3%) | 721<br>(94.6%) | 8 (1.0%) | 80 (10.5%) | 83 (10.9%) | 599 (78.6%) | 590 |
| Total *in situ* |  | 6095 | 79<br>(1.3%) | 5734<br>(94.1%) | 282<br>(4.6%) | 325 (5.3%) | 2825<br>(46.3%) | 2945<br>(48.3%) | 2127 |

**Table S3.** Tether-influence and activity classification results for individual jellyfish.

| Deployment ID | Representative pulse frequency (pulses/sec) | Unannotated data* classified as influenced | Unannotated data* classified as drifting (out of time classified as uninfluenced) |
| --- | --- | --- | --- |
| S1-1 | 0.260 | 35.1% | 19.1% |
| S1-2 | 0.487 | 21.2% | 0% |
| S1-3 | 0.615 | 6.0% | 0.9% |
| S2-1 | 0.520 | 3.3% | 0% |
| S2-2 | 0.466 | 5.0% | 0.1% |
| S3-1 | 0.275 | 28.2% | 5.6% |
| S3-2 | 0.422 | 9.8% | 0.6% |
| S3-3 | 0.380 | 11.4% | 2.7% |

\* Rightmost column of Table S2.

**Movie S1. Examples of annotated *in situ* and laboratory footage**. In order, uninfluenced *in situ* swimming, tether-influenced *in situ* swimming, *in situ* swimming with unknown tether status, uninfluenced *in situ* drifting, tether-influenced *in situ* drifting, and swimming and drifting in the MBARI Test Tank.

## References

Ambroise, C. and McLachlan, G. J. (2002). Selection bias in gene extraction on the basis of microarray gene-expression data. Proc. Natl. Acad. Sci. U. S. A. 99, 6562–6566.

Båmstedt, U., Kaartvedt, S. and Youngbluth, M. (2003). An evaluation of acoustic and video methods to estimate the abundance and vertical distribution of jellyfish. J. Plankton Res. 25, 1307–1318.

Block, B. A., Jonsen, I. D., Jorgensen, S. J., Winship, A. J., Shaffer, S. A., Bograd, S. J., Hazen, E. L., Foley, D. G., Breed, G. A., Harrison, A.-L., et al. (2011). Tracking apex marine predator movements in a dynamic ocean. Nature 475, 86–90.

Blockeel, H., Kersting, K., Nijssen, S. and Železný, F. eds. (2013). Machine Learning and Knowledge Discovery in Databases: European Conference, ECML PKDD 2013, Prague, Czech Republic, September 23-27, 2013, Proceedings, Part III. Springer, Berlin, Heidelberg.

Boyd, K., Eng, K. H. and Page, C. D. (2013). Area under the Precision-Recall Curve: Point Estimates and Confidence Intervals. In Machine Learning and Knowledge Discovery in Databases, pp. 451–466. Springer Berlin Heidelberg.

Brewster, L. R., Dale, J. J., Guttridge, T. L., Gruber, S. H., Hansell, A. C., Elliott, M., Cowx, I. G., Whitney, N. M. and Gleiss, A. C. (2018). Development and application of a machine learning algorithm for classification of elasmobranch behaviour from accelerometry data. Mar. Biol. 165, 62.

Brown, D. D., Kays, R., Wikelski, M., Wilson, R. and Klimley, A. P. (2013). Observing the unwatchable through acceleration logging of animal behavior. Animal Biotelemetry 1, 20.

Bunescu, R., Ge, R., Kate, R. J., Marcotte, E. M., Mooney, R. J., Ramani, A. K. and Wong, Y. W. (2005). Comparative experiments on learning information extractors for proteins and their interactions. Artif. Intell. Med. 33, 139–155.

Carroll, G., Slip, D., Jonsen, I. and Harcourt, R. (2014). Supervised accelerometry analysis can identify prey capture by penguins at sea. J. Exp. Biol. 217, 4295–4302.

Cawley, G. C. and Talbot, N. L. C. (2010). On Over-fitting in Model Selection and Subsequent Selection Bias in Performance Evaluation. J. Mach. Learn. Res. 11, 2079–2107.

Colin, S. P. and Costello, J. H. (2002). Morphology, swimming performance and propulsive mode of six co-occurring hydromedusae. J. Exp. Biol. 205, 427–437.

Colin, S. P., Costello, J. H. and Klos, E. (2003). In situ swimming and feeding behavior of eight co-occurring hydromedusae. Mar. Ecol. Prog. Ser. 253, 305–309.

Costello, J. H., Klos, E. and Ford (1998). In situ time budgets of the scyphomedusae *Aurelia aurita*, *Cyanea* sp., and *Chrysaora quinquecirrha*. J. Plankton Res. 20, 383–391.

Dabiri, J. O., Colin, S. P., Katija, K. and Costello, J. H. (2010). A wake-based correlate of swimming performance and foraging behavior in seven co-occurring jellyfish species. J. Exp. Biol. 213, 1217–1225.

Dash, M. and Liu, H. (1997). Feature selection for classification. Intelligent Data Analysis 1, 131–156.

Davis, J. and Goadrich, M. (2006). The Relationship Between Precision-Recall and ROC Curves. In Proceedings of the 23rd International Conference on Machine Learning, pp. 233–240. New York, NY, USA: ACM.

Diebel, J. (2006). Representing attitude: Euler angles, unit quaternions, and rotation vectors. Matrix 58, 1–35.

Duarte, M. (2013). Notes on Scientific Computing for Biomechanics and Motor Control. Github.

Duda, R. O., Hart, P. E. and Stork, D. G. (2000). Pattern Classification (2Nd Edition). New York, NY, USA: Wiley-Interscience.

Fisher, R. A. (1936). The use of multiple measurements in taxonomic problems. Ann. Eugen. 7, 179–188.

Fawcett, T. (2006). An introduction to ROC analysis. Pattern Recognit. Lett. 27, 861–874.

Fossette, S., Gleiss, A. C., Chalumeau, J., Bastian, T., Armstrong, C. D., Vandenabeele, S., Karpytchev, M. and Hays, G. C. (2015). Current-oriented swimming by jellyfish and its role in bloom maintenance. Curr. Biol. 25, 342–347.

Fossette, S., Katija, K., Goldbogen, J. A., Bograd, S., Patry, W., Howard, M. J., Knowles, T., Haddock, S. H. D., Bedell, L., Hazen, E. L., et al. (2016). How to tag a jellyfish? A methodological review and guidelines to successful jellyfish tagging. J. Plankton Res. 38, 1347–1363.

Gemmell, B. J., Colin, S. P. and Costello, J. H. (2018). Widespread utilization of passive energy recapture in swimming medusae. J. Exp. Biol. 221.

Gleiss, A. C., Wilson, R. P. and Shepard, E. L. C. (2011). Making overall dynamic body acceleration work: on the theory of acceleration as a proxy for energy expenditure: Acceleration as a proxy for energy expenditure. Methods Ecol. Evol. 2, 23–33.

Goldbogen, J. A., Calambokidis, J., Shadwick, R. E., Oleson, E. M., McDonald, M. A. and Hildebrand, J. A. (2006). Kinematics of foraging dives and lunge-feeding in fin whales. J. Exp. Biol. 209, 1231–1244.

Guyon, I. and Elisseeff, A. (2003). An Introduction to Variable and Feature Selection. J. Mach. Learn. Res. 3, 1157–1182.

Halsey, L. G., Green, J. A., Wilson, R. P. and Frappell, P. B. (2009). Accelerometry to Estimate Energy Expenditure during Activity: Best Practice with Data Loggers. Physiol. Biochem. Zool. 82, 396–404.

Hastie, T., Tibshirani, R. and Friedman, J. (2009). The Elements of Statistical Learning: Data Mining, Inference, and Prediction. Springer Science & Business Media.

Hays, G. C., Doyle, T. K., Houghton, J. D. R., Lilley, M. K. S., Metcalfe, J. D. and Righton, D. (2008). Diving behaviour of jellyfish equipped with electronic tags. J. Plankton Res. 30, 325–331.

Hays Graeme C., Bastian Thomas, Doyle Thomas K., Fossette Sabrina, Gleiss Adrian C., Gravenor Michael B., Hobson Victoria J., Humphries Nicolas E., Lilley Martin K. S., Pade Nicolas G., et al. (2012). High activity and Lévy searches: jellyfish can search the water column like fish. Proceedings of the Royal Society B: Biological Sciences 279, 465–473.

Hays, G. C., Ferreira, L. C., Sequeira, A. M. M., Meekan, M. G., Duarte, C. M., Bailey, H., Bailleul, F., Bowen, W. D., Caley, M. J., Costa, D. P., et al. (2016). Key Questions in Marine Megafauna Movement Ecology. Trends Ecol. Evol. 31, 463–475.

Honda, N., Watanabe, T. and Matsushita, Y. (2009). Swimming depths of the giant jellyfish Nemopilema nomurai investigated using pop-up archival transmitting tags and ultrasonic pingers. Fish. Sci. 75, 947–956.

Hurley, N. and Rickard, S. (2009). Comparing Measures of Sparsity. IEEE Trans. Inf. Theory 55, 4723–4741.

Jeantet, L., Dell’Amico, F., Forin-Wiart, M.-A., Coutant, M., Bonola, M., Etienne, D., Gresser, J., Regis, S., Lecerf, N., Lefebvre, F., et al. (2018). Combined use of two supervised learning algorithms to model sea turtle behaviours from tri-axial acceleration data. J. Exp. Biol. 221.

Johnson, M. P. and Tyack, P. L. (2003). A digital acoustic recording tag for measuring the response of wild marine mammals to sound. IEEE J. Oceanic Eng. 28, 3–12.

Kaartvedt, S., Klevjer, T. A., Torgersen, T., Sørnes, T. A. and Røstad, A. (2007). Diel vertical migration of individual jellyfish (Periphylla periphylla). Limnol. Oceanogr. 52, 975–983.

Kaartvedt, S., Ugland, K. I., Klevjer, T. A., Røstad, A., Titelman, J. and Solberg, I. (2015). Social behaviour in mesopelagic jellyfish. Sci. Rep. 5, 11310.

Klevjer, T. A., Kaartvedt, S. and Båmstedt, U. (2009). In situ behaviour and acoustic properties of the deep living jellyfish Periphylla periphylla. J. Plankton Res. 31, 793–803.

Kohavi, R. (1995). A Study of Cross-validation and Bootstrap for Accuracy Estimation and Model Selection. In Proceedings of the 14th International Joint Conference on Artificial Intelligence - Volume 2, pp. 1137–1143. San Francisco, CA, USA: Morgan Kaufmann Publishers Inc.

Kooyman, G. L. (2004). Genesis and evolution of bio-logging devices: 1963-2002. Mem. Natl Inst. Polar Res., Spec. Issue 58, 15–22.

Ladds, M. A., Thompson, A. P., Slip, D. J., Hocking, D. P. and Harcourt, R. G. (2016). Seeing It All: Evaluating Supervised Machine Learning Methods for the Classification of Diverse Otariid Behaviours. PLoS One 11, e0166898.

Liu, H. and Motoda, H. (1998). Feature Selection for Knowledge Discovery and Data Mining. Springer Science & Business Media.

MacKay, D. J. C. and Mac, D. J. (2003). Information Theory, Inference and Learning Algorithms. Cambridge University Press.

Manning, C. D. and Schütze, H. (1999). Foundations of Statistical Natural Language Processing. MIT Press.

Martín López, L. M., Aguilar de Soto, N., Miller, P. and Johnson, M. (2016). Tracking the kinematics of caudal-oscillatory swimming: a comparison of two on-animal sensing methods. J. Exp. Biol. 219, 2103–2109.

Matanoski, J., Hood, R. and Purcell, J. (2001). Characterizing the effect of prey on swimming and feeding efficiency of the scyphomedusa Chrysaora quinquecirrha. Mar. Biol. 139, 191–200.

Mills, C. E. (1984). Density is altered in hydromedusae and ctenophores in response to changes in salinity. Biol. Bull. 166, 206–215.

Mooney, T. A., Katija, K., Shorter, K. A., Hurst, T., Fontes, J. and Afonso, P. (2015). ITAG: an eco-sensor for fine-scale behavioral measurements of soft-bodied marine invertebrates. Animal Biotelemetry 3, 31.

Moriarty, P. E., Andrews, K. S., Harvey, C. J. and Kawase, M. (2012). Vertical and horizontal movement patterns of scyphozoan jellyfish in a fjord-like estuary. Mar. Ecol. Prog. Ser. 455, 1–12.

Pan, S. J. and Yang, Q. (2010). A Survey on Transfer Learning. IEEE Trans. Knowl. Data Eng. 22, 1345–1359.

Purcell, J. E. (2009). Extension of methods for jellyfish and ctenophore trophic ecology to large-scale research. Hydrobiologia 616, 23–50.

Rasmussen, K., Palacios, D. M., Calambokidis, J., Saborío, M. T., Dalla Rosa, L., Secchi, E. R., Steiger, G. H., Allen, J. M. and Stone, G. S. (2007). Southern Hemisphere humpback whales wintering off Central America: insights from water temperature into the longest mammalian migration. Biol. Lett. 3, 302–305.

Reunanen, J. (2003). Overfitting in Making Comparisons Between Variable Selection Methods. J. Mach. Learn. Res. 3, 1371–1382.

Richardson, M. and Domingos, P. (2006). Markov logic networks. Mach. Learn. 62, 107–136.

Rife, J. and Rock, S. M. (2003). Segmentation methods for visual tracking of deep-ocean jellyfish using a conventional camera. IEEE J. Oceanic Eng. 28, 595–608.

Rutz, C. and Hays, G. C. (2009). New frontiers in biologging science. Biol. Lett. 5, 289–292.

Sato, K., Mitani, Y., Cameron, M. F., Siniff, D. B. and Naito, Y. (2003). Factors affecting stroking patterns and body angle in diving Weddell seals under natural conditions. J. Exp. Biol. 206, 1461–1470.

Sequeira, A. M. M., Rodríguez, J. P., Eguíluz, V. M., Harcourt, R., Hindell, M., Sims, D. W., Duarte, C. M., Costa, D. P., Fernández-Gracia, J., Ferreira, L. C., et al. (2018). Convergence of marine megafauna movement patterns in coastal and open oceans. Proc. Natl. Acad. Sci. U. S. A. 115, 3072–3077.

Seymour, J. E., Carrette, T. J. and Sutherland, P. A. (2004). Do box jellyfish sleep at night? Med. J. Aust. 181, 707.

Shepard, E. L. C., Wilson, R. P., Halsey, L. G., Quintana, F., Gómez Laich, A., Gleiss, A. C., Liebsch, N., Myers, A. E. and Norman, B. (2008). Derivation of body motion via appropriate smoothing of acceleration data. Aquat. Biol. 4, 235–241.

Sims, D. W., Southall, E. J., Humphries, N. E., Hays, G. C., Bradshaw, C. J. A., Pitchford, J. W., James, A., Ahmed, M. Z., Brierley, A. S., Hindell, M. A., et al. (2008). Scaling laws of marine predator search behaviour. Nature 451, 1098–1102.

Smialowski, P., Frishman, D. and Kramer, S. (2010). Pitfalls of supervised feature selection. Bioinformatics 26, 440–443.

Sugiyama, M., Krauledat, M. and Müller, K.-R. (2007). Covariate Shift Adaptation by Importance Weighted Cross Validation. J. Mach. Learn. Res. 8, 985–1005.

Varma, S. and Simon, R. (2006). Bias in error estimation when using cross-validation for model selection. BMC Bioinformatics 7, 91.

Watanabe, Y. Y. and Takahashi, A. (2013). Linking animal-borne video to accelerometers reveals prey capture variability. Proc. Natl. Acad. Sci. U. S. A. 110, 2199–2204.

Weise, M. J., Harvey, J. T. and Costa, D. P. (2010). The role of body size in individual-based foraging strategies of a top marine predator. Ecology 91, 1004–1015.

Whitney, A. W. (1971). A Direct Method of Nonparametric Measurement Selection. IEEE Trans. Comput. C-20, 1100–1103.

Wilson, R. P., White, C. R., Quintana, F., Halsey, L. G., Liebsch, N., Martin, G. R. and Butler, P. J. (2006). Moving towards acceleration for estimates of activity-specific metabolic rate in free-living animals: the case of the cormorant. J. Anim. Ecol. 75, 1081–1090.

Zhang, K., Schölkopf, B., Muandet, K. and Wang, Z. (2013). Domain Adaptation under Target and Conditional Shift. In International Conference on Machine Learning, pp. 819–827.

Zonoobi, D., Kassim, A. A. and Venkatesh, Y. V. (2011). Gini Index as Sparsity Measure for Signal Reconstruction from Compressive Samples. IEEE J. Sel. Top. Signal Process. 5, 927–932.

